# Backtranslation of human RNA biosignatures of tuberculosis disease risk into the preclinical pipeline is condition dependent

**DOI:** 10.1101/2024.06.21.600067

**Authors:** Hannah Painter, Sasha E. Larsen, Brittany D. Williams, Hazem F. M. Abdelaal, Susan L. Baldwin, Helen A. Fletcher, Andrew Fiore-Gartland, Rhea N. Coler

## Abstract

It is not clear whether human progression to active tuberculosis disease (TB) risk signatures are viable endpoint criteria for evaluations of treatments in clinical or preclinical development. TB is the deadliest infectious disease globally and more efficacious vaccines are needed to reduce this mortality. However, the immune correlates of protection for either preventing infection with *Mycobacterium tuberculosis* or preventing TB disease have yet to be completely defined, making the advancement of candidate vaccines through the pipeline slow, costly, and fraught with risk. Human-derived correlate of risk (COR) gene signatures, which identify an individual’s risk to progressing to active TB disease, provide an opportunity for evaluating new therapies for TB with clear and defined endpoints. Though prospective clinical trials with longitudinal sampling are prohibitively expensive, characterization of COR gene signatures is practical with preclinical models. Using a 3Rs (Replacement, Reduction and Refinement) approach we reanalyzed heterogeneous publicly available transcriptional datasets to determine whether a specific set of COR signatures are viable endpoints in the preclinical pipeline. We selected RISK6, Sweeney3 and BATF2 human-derived blood-based RNA biosignatures because they require relatively few genes to assign a score and have been carefully evaluated across several clinical cohorts. Excitingly, these data provide proof-of-concept that human COR signatures seem to have high fidelity across several tissue types in the preclinical TB model pipeline and show best performance when the model most closely reflected human infection or disease conditions. Human-derived COR signatures offer an opportunity for high-throughput preclinical endpoint criteria of vaccine and drug therapy evaluations.

**One Sentence Summary:** Human-derived biosignatures of tuberculosis disease progression were evaluated for their predictive fidelity across preclinical species and derived tissues using available public data sets.

## INTRODUCTION

Tuberculosis disease (TB), caused by the bacterial pathogen *Mycobacterium tuberculosis* (M.tb), was the leading infectious killer globally for the four years (*1*) predating the SARS-CoV-2 pandemic and was again the widespread frontrunner from 2022 (*2*). Approximately 1.3 million deaths, including over 150,000 people living with HIV, in 2023 worldwide were attributed to TB (*3*). The World Health Organization (WHO) estimates that COVID-19 related disruptions in care could result in an additional half million deaths from TB (*4*). The burden of TB continues to disproportionately affect low and middle income countries (*5*). While the classic infection risks for M.tb are fairly well known and studied (*6–10*), the majority of individuals who harbor M.tb do not advance to active TB disease (*11–15*). Less well known are the factors that influence an individual’s progression through the spectrum of TB disease (*14, 16, 17*), or what determines their risk of advancing to an active, transmissible state. Indeed, the current methods to identify M.tb infection, the tuberculin skin test (TST) and interferon-γ release assay (IGRA), have poor specificity for TB disease in endemic populations, including for those individuals living with HIV (*18, 19*). The TB field is in need of better triage criteria that can allow targeted identification of active and incipient subclinical TB disease.

Highly specific predictive correlate of risk (COR) biomarkers or diagnostics could enable risk-stratified preventive interventions. These tools may avoid unnecessary treatment of people who would likely remain healthy, while also interrupting incipient TB disease in infected adults, leading to a reduction in morbidity and transmission. Studies are ongoing to evaluate several clinical applications (*20–23*). Human peripheral blood-based biosignatures have been able to selectively identify TB disease from other infections, distinguish latent TB infection (LTBI) and predict progression to active TB disease (*24–29*). Correlative biomarkers for TB-related disease states are well reviewed here (*30–32*). Iterative large-scale human clinical trials to examine the performance of multiple host biomarkers would be cost prohibitive, however assessing human-derived RNA risk signatures of progression to active disease in the preclinical pipeline may afford rigor and fidelity to select efficacy endpoints used to screen products and accelerate preclinical development.

Understanding the strengths or limitations of back-translating human-derived COR RNA signatures into the preclinical pipeline may help streamline down-selection of therapeutic vaccine and drug candidates and better align preclinical models with proposed clinical trial efficacy endpoints. However, to date, few published studies have evaluated the appropriateness of leveraging human-derived RNA COR gene signatures, including RISK6, Sweeney3 or BATF2, in preclinical models (*33*). Given the abundance of host-derived gene expression data published, in this work we aggregated a cascade of existing public transcriptomic datasets across the spectrum of preclinical TB disease models (mouse, guinea pig [GP], rabbit and non-human primates [NHP]) and interrogated them for expression of select COR gene signatures and risk score correlation with primary disease endpoints. Our primary objective was to determine if human-derived RNA gene signatures predicting progression to active TB disease could be back-translated into animal models. Our secondary objective was to identify whether certain experimental conditions, such as infectious dose, were limitations for RNA COR signature use. We hypothesized that along with bacterial burden, pulmonary inflammation-induced damage may correlate with increasing COR gene signatures across preclinical models. Further, we expected to identify models of vaccine and drug interventions that lower COR gene expression.

## RESULTS

Parsimonious (1–6 genes) COR biosignatures, RISK6 (*27*), Sweeney3 (*34*) and BATF2 (*35*) were selected for use in these analyses; this increased the chance of all genes/homologs being present in data sets analyzed (**Table 1**). Biosignatures with a high number of genes may lose sensitivity and the ability to compare between data sets if genes are present in some but not others. In published human cohort studies, a higher number of genes was not found to correlate with better performance in a head-to-head comparison of 16 different biosignatures (*36*). The three biosignatures chosen for this work were included in two critical meta-analyses, which used independent validation cohorts to demonstrate that sensitivity and specificity remain high, particularly when in close proximity to TB disease status (*32, 37*). Interestingly, many genes contributing to each signature score are present across several gene signatures developed in human clinical studies (see Supplemental Table 2 Gene Matrix here (*37*)) and have direct immune regulatory functions (**Supplemental Table 1**). Further, Sweeney3 and RISK6 are under development as point-of-care tests and have been shown to correlate with pathology scores in humans (*27, 38, 39*). Pathology scores can be readily assessed in the majority of preclinical animal models of TB, and a number of the gene expression datasets reanalyzed in this study included pathology scores.

**Table 1.** Description of down-selected human TB transcriptional risk signatures.

| Name | Gene composition | Score equation | Discovery cohort | Current applications |
| --- | --- | --- | --- | --- |
| RISK6 (27) | GBP2, FCGR1B*, SERPING1, TUBGCP6, TRMT2A, SDR39U1 | geometric mean(TUBGCP6, TRMT2A, SDR39U1) – geometric mean(GBP2, FCGR1B, SERPING1) | HIV-negative South African adolescents | TB disease risk, diagnosis and treatment response |
| Sweeney3 (34) | GBP5, DUSP3, KLF2 | mean(GBP5 + DUSP3) – KLF2 | HIV-positive and –negative adults, various sites | TB disease risk, diagnosis, disease severity and treatment response |
| BATF2 (35) | BATF2 | BATF2 | HIV-negative UK adults | TB disease risk and diagnosis; differentiation between pulmonary and extra-pulmonary TB |
| *In all murine datasets, the mouse ortholog Fcgr1 was used; NHP Ahmed, FCGR1; Gideon, FCGR1B; Hansen, FCGR1A. Otherwise gene homologs were used for each species. |  |  |  |  |

This reanalysis of published data provides proof-of-concept that genes used in select human COR signatures also predict progression to active TB disease in preclinical models of M.tb infection in the context of the spectrum of TB disease (naïve, latent and active), as well as identifying drug treatment success. These signatures may be suitable for use as surrogate vaccine intervention endpoints with appropriate kinetic considerations.

### Primary data discovery and inclusion-exclusion criteria

Searches for primary data for inclusion in this reanalysis work were executed through Gene Expression Omnibus (GEO) repository as well as PubMed. We chose to include *in vivo* preclinical models of TB experimental infection including mice, GPs, rabbits and NHPs to cover a spectrum of disease outcomes and intervention strategies. GEO entries were collated with the following search terms “mus tuberculosis AND Mus musculus” (69 total hits), “macaca tuberculosis” (4 total hits), “oryctolagus tuberculosis” (7 total hits) and “cavia tuberculosis” (1 hit). Since some data is presented in supplemental figures and not directly stored in GEO, we executed tandem searches in PubMed using combinations of the following terms: tuberculosis, M.tb, biomarkers, RNA, risk scores, preclinical models, mouse, NHP, guinea pig, rabbit, challenge, microarray, sequencing, host gene expression, vaccine, or drug treatment. Four non-redundant data sets were identified on PubMed that were not present in results from GEO searches. The aim of this work is to evaluate whether these three selected biosignatures can be readily applied across the preclinical pipeline. We therefore omitted data involving cell lines or sorted primary cells, or single-cell RNA-seq data, since these source tissues add complexity to analysis, are more mechanistic and would reduce throughput for the pipeline (**Table 2**). Inclusion criteria for gene expression data for the reanalysis involved was: data derived from non-review sources; data generated from tissue within a preclinical *in vivo* experimental infection study; study group size within the dataset must be ≥2; and dataset contains at least one of the three selected biosignatures (**Table 2**). While datasets with n=2 were collected, they were used for observational trends only and not assessed statistically.

**Table 2.** Selection of data for analysis.

| Inclusion criteria | Exclusion criteria |
| --- | --- |
| Contains primary data, non-review based | Single-cell RNA-seq |
| Contains complete gene set in at least one biosignature | Cell lines |
| Based on tissue derived from *in vivo* preclinical models of TB experimental infection | Sorted primary cell subsets |
| Group size $n \geq 2$ | |

Through GEO and PubMed we identified a total of 30 sources of primary gene expression data meeting our inclusion and exclusion criteria which were further reanalyzed for this work (**Tables 3-5**). Datasets identified but excluded can be found in **Supplemental Table 2** with rationale for exclusion. These sources provided 27 *Mus musculus* (mouse) and 3 *Macaca* (NHP) data sets. Data selected for reanalysis represents a spectrum of models with different experimental infection doses (**Table 3**), kinetics with respect to time since infection (**Table 4**), tissues of interest, as well as vaccine and chemotherapeutic interventions (**Table 5**). Throughout we consistently use the primary authors’ definition of infection dose as “high” or “low” and state CFU range as reported in materials and methods.

**Table 3.** Datasets involving different infection doses of M.tb.

| GEO (ref) | First author; year | Sample | Species | Sex | Infection dose | M.tb strain |
| --- | --- | --- | --- | --- | --- | --- |
| GSE158807<br>(33) | Plumlee; 2020 | WB | *Mus musculus*;<br>C57Bl/6 | F | Ultra-low: 1-3 CFU;<br>conventional: 50-100 CFU | H37Rv |
| n/a<br>(41) | Ahmed; 2020 | Lung | *Macaca mulatta* | M & F | Low: 10 CFU;<br>high: 100 CFU | CDC1551 |
| GSE179417<br>(42) | Koyuncu;<br>2021 | Lung | *Mus musculus*;<br>D.O. | F | 25-100 CFU | Erdman |
| n/a<br>(41) | Ahmed; 2020 | Lung | *Mus musculus*;<br>D.O. | M & F | 100 CFU | HN878 |
| GSE137093<br>(43) | Moreira-Teixeira;<br>2019 | Lung | *Mus musculus*;<br>C57Bl/6 &<br>C3HeB/FeJ | F | At day 3: low: 100-450 CFU; high: 700-900 | H37Rv &<br>HN878 |
| GSE137092<br>(43) | Moreira-Teixeira;<br>2019 | WB<sup>#</sup> | *Mus musculus*;<br>C57Bl/6 &<br>C3HeB/FeJ | F | At day 3: low: 100-450 CFU; high: 700-900 | H37Rv &<br>HN878 |
| D.O.: diversity outbred mice; WB: whole blood sample; M.tb infection is via aerosol unless otherwise noted. |  |  |  |  |  |  |

**Table 4.** Datasets involving kinetic timepoints post infection.

| GEO (ref) | First author; year | Sample | Species | Sex | Infection dose | M.tb strain | Sampling days post infection |
| --- | --- | --- | --- | --- | --- | --- | --- |
| GSE124688 (44) | Ault; 2019 | WB | *Mus musculus*; C57Bl/6 | F | 50-100 CFU | Erdman | 30, 60, 90, 120 and 150 |
| GSE21149 (45) | Aranday-Cortes; 2011 | Lung, spleen | *Mus musculus*; BALB/c | F | 600 CFU | M. bovis (i.n.) | 0, 3 and 14 |
| GSE140944 (43) | Moreira-Teixeira; 2019 | Lung | *Mus musculus*; BALB/c | F | 100-1000 CFU | H37Rv | 0, 14, 21, 28, 57 and 129 |
| GSE140943 (43) | Moreira-Teixeira; 2019 | Blood | *Mus musculus*; BALB/c | F | 100-1000 CFU | H37Rv | 0, 14, 21, 28, 56 and 138 |
| GSE89389 (46) | Domaszewska; 2017 | WB | *Mus musculus*; C57Bl/6 & 129S2 | M & F | “low dose” | H37Rv | 0, 1, 7, 14 and 21 |
| GSE64045 (47) | Mehra; 2014 | Lung | *Mus musculus*; C3HeB/FeJ | M | ~100 CFU | H37Rv | 28, 56 and 112 |
| GSE23014 (48) | Kang; 2010 | Lung | *Mus musculus*; C57Bl/6 | M & F | 100 CFU | H37Rv | 0, 12, 15 and 21 |
| GSE84152 (49) | Gideon; 2016 | WB | *Macaca fascicularis* | M & F | 25 CFU | Erdman (instill.) | 0, 1, 3, 7, 10, 20, 30, 42, 56, |
|  |  |  |  |  |  |  | 90, 120, 150 and 180 |
| GSE169541<br>(50) | Cerezo-Cortés;<br>2021 | Lung | *Mus musculus*;<br>BALB/c | M | 250 CFU | Beijing-like<br>(instill.) | 3, 14, 28 and 60 |
| GSE168486<br>(51) | Naqvi;<br>2021 | Lung | *Mus musculus*;<br>C57Bl/6 &<br>MGL-1<sup>-/-</sup> | M & F | 100 CFU | H37Rv<br>(i.n.) | 14 and 56 |
| M.tb infection is via aerosol unless otherwise noted; i.n. = intranasal infection; instill = intratracheal or intrabronchial instillation, WB = whole blood. |  |  |  |  |  |  |  |

**Table 5.** Datasets involving anti-M.tb interventions.

| GEO (ref) | First author; year | Sample | Species | Sex | Infection dose | M.tb Strain | Intervention |
| --- | --- | --- | --- | --- | --- | --- | --- |
| GSE64167 (52) | Wilk; 2013 | Lung | *Mus musculus*; A/Sn & I/St | M | 5 x 10<sup>6</sup> CFU | H37Rv (i.v.) | Genetic |
| GSE166114 (53) | Ji; 2021 | Lung | *Mus musculus*; C57Bl/6 & Sp110 & Sp140 | M & F | 20-50 CFU | Erdman | Genetic |
| GSE165871 (54) | Bohrer; 2021 | Lung | *Mus musculus*; C57Bl/6 & ΔdblGata | M & F | Low:100-300<br>High: 1000-1500 CFU | H37Rv | Genetic |
| GSE141207 (55) | Moreira-Teixeira; 2019 | Blood and lung | *Mus musculus*; C57Bl/6 | F | 200 CFU | HN878 | α-GM-CSF |
| GSE55183 (56) | Dutta; 2013 | Lung | *Mus musculus*; C3HeB/FeJ | F | “OD600 approx 1.0” | H37Rv | α-TNF |
| GSE127263 (57) | Matarese; 2019 | Spleen | *Mus musculus*; DBA/2 | F | 10<sup>5</sup> CFU | H37Rv (i.v.) | Caloric restriction |
| GSE57275 (58) | Singal; 2014 | Lung | *Mus musculus*; C57Bl/6 | F | 10<sup>3</sup> CFU | H37Rv | Metformin |
| GSE179437 (59) | Gandotra; 2021 | Lung | *Mus musculus*; C3HeB/FeJ | M & F | Low: 100<br>High: 500 CFU | Erdman | T863 |
| GSE99625 (60) | Dutta; 2017 | Lung | *Mus musculus*; BALB/c | F | 5 x 10<sup>3</sup> CFU | H37Rv | RHZE |
| GSE83188<br>(61) | Subbian; 2016 | Lung | *Mus musculus*;<br>B6D2F1 | F | 100-150<br>CFU | CDC1551 | CC 11050 |
| GSE48027<br>(62) | Manca;<br>2013 | Lung | *Mus musculus*;<br>B6D2F1 | F | 2.5 log<sub>10</sub> | CDC1551 | PZA |
| GSE44825<br>(63) | Rodrigues;<br>2013 | Lung | *Mus musculus*;<br>BALB/c | F | 10<sup>5</sup> CFU | H37Rv<br>(instill.) | RH |
| GSE97835<br>(64) | Park;<br>2017 | Lung | *Mus musculus*;<br>C57Bl/6 | n/a | 200-300<br>CFU | Erdman | HZE |
| GSE102440<br>(65) | Hansen;<br>2017 | WB | *Macaca mulatta* | M | 10-25<br>CFU | Erdman<br>(instill.) | Vaccination |
| M.tb infection is via aerosol unless otherwise noted; E = ethambutol; GM-CSF = granulocyte-macrophage colony-stimulating factor; H = isoniazid; i.v. = intravenous infection; instill = intratracheal or intrabronchial instillation; R = rifampin; TNF = tumor necrosis factor; WB = whole blood; Z/PZA = pyrazinamide. |  |  |  |  |  |  |  |

### Dose

Refinement of preclinical aerosol infection models has allowed TB researchers to consistently tailor the infection doses of M.tb from as few as one bacillus to several thousand (*33, 40*). These developments better represent the understanding of natural human exposure to M.tb where few or one bacilli can establish infection. Since host COR signatures have been demonstrated to aid prediction of progression to active disease, often corresponding with loss of control of M.tb in granulomas and expansion of the bacillary load, we hypothesized that infection dose may positively correlate with COR signature expression in these data sets. This reanalysis encompassed six datasets meeting our inclusion/exclusion criteria which included NHP and mouse (diversity outbred [D.O.], C57BL/6 and C3HeB/FeJ) samples derived from lungs and whole blood (WB), across several M.tb strains including M.tb HN878, H37Rv, CDC1551 and Erdman (**Table 3**).

### M.tb infection dose modifies resolution of bacterial burden and signature score expression

The Plumlee *et al* (GSE158807) dataset contained the lowest infection dose and a higher, conventional dose (CD) (*33*). We compared risk signature scores in WB isolated from naïve mice, mice by infected with a low-dose aerosol (LDA, 50-100 CFU, also termed CD by the authors) and ultra-low dose aerosol (ULDA, 1-3 CFU) infected mice. Here we were interested to know whether mouse-level variability in signature expression correlated with dose at time of infection and/or resulting CFU at designated endpoints. In general, each score was able to differentiate CD infection from either naïve mice or ULDA mice (**Figure 1**). Both CD and early ULDA Risk6 scores were higher than naïve, with a general increase in score between ULDA and CD although this was not significant (**Figure 1 A**). Sweeney3 scores between cohorts were able to differentiate significant differences between each group, again where CD scores were higher than ULDA (**Figure 1 B**). Interestingly BATF2 scores were very similar between all groups tested and the only difference observed was between late-isolated ULDA and CD (**Figure 1 C**). This study had corresponding lung CFU meta-data that could be compared with scores between groups as well. When comparing lung CFU values to each individual animal score we observe that Risk6 (**Figure 1 D**), Sweeney3 (**Figure 1 E**) and BATF2 (**Figure 1 F**) scores each correlate with pulmonary bacterial burden. It is difficult to disentangle whether these relationships between signature expression and outcome CFU are singular or also involve early innate responses to infection not evaluated here. Overall Risk6 variability correlated strongest with outcome CFU (R = 0.5011, p < 0.0001) independent of infection treatment group, observed in **Figure 1 B** where group colors overlap. Similarly, for BATF2, which correlates with outcome even though it is equal among the challenge doses, suggesting it has more to do with outcome than initial challenge. Whereas, Sweeney3 scores seem to separate more strongly by infection treatment group (**Figure 1 D**).

**Figure 1.**
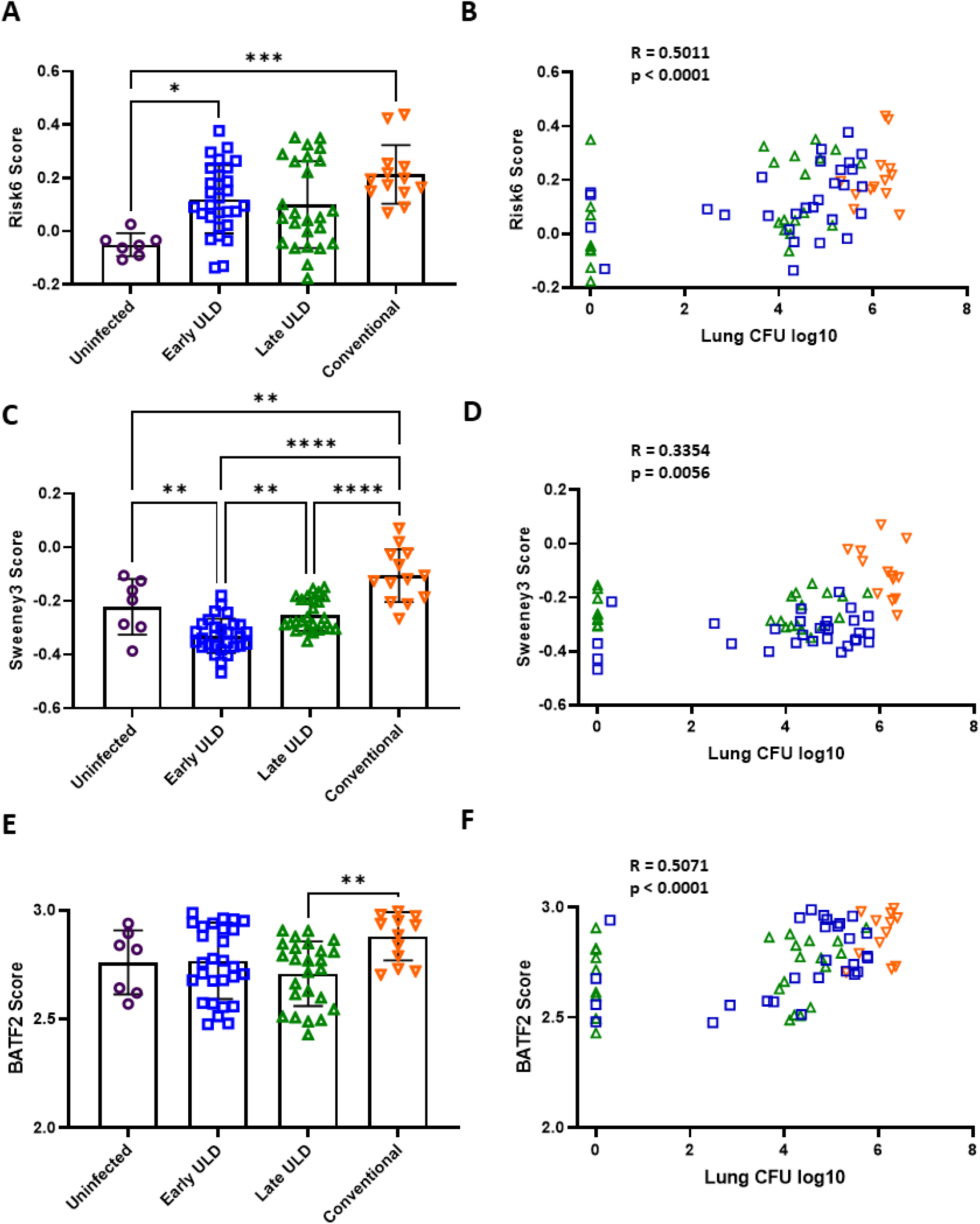
Risk signature scores by infection dose in a mouse model. In Plumlee *et al* (GSE158807) C57BL/6 mice were infected with a CD/LDA (50-100 CFU) or ULDA (1-3 CFU) of M.tb H37Rv and host RNA was collected from WB at 24-days post infection. Samples were analyzed using Agilent-074809 SurePrint G3 Mouse GE v2 8x60K Microarray and compared with naïve (open purple circles) at early (ULDA only, open blue squares) and late timepoints (CD: open orange upside-down triangle and ULDA: open green triangle). **A**) Risk6, **C**) Sweeney3 and **E**) BATF2 scores were calculated for each mouse and cohorts were compared using one-way ANOVA with Tukey’s multiple comparison test correction. Significant comparisons are denoted by asterisks in the figure where * = p ≤ 0.05, ** = p ≤ 0.01, *** = p ≤ 0.001, **** = p ≤ 0.0001. Log_10_ lung CFU values were matched for each animal to **B**) Risk6, **D**) Sweeney3 and **F**) BATF2 scores for correlation using a Spearman rank-based correlation test. R and p values are shown for each test in the figure.

### Infection dose stratifies controller vs progressor scores better than pathology or morbidity endpoints

Three studies categorized cohorts as “controllers” or “progressors” after different infection doses of M.tb and are presented here together. In Ahmed *et al* (*41*) Indian rhesus macaques were infected with either 10 CFU M.tb CDC1551 and lung tissue was collected at 5-8 weeks post infection (controllers), or 100 CFU M.tb CDC1551 and samples collected 22-24 weeks post infection (progressors). Gene expression analysis was performed by RNA-seq. Koyuncu and colleagues infected D.O. mice with 25-100 CFU M.tb Erdman and lung RNA was analyzed by Affymetrix Mouse Gene 2.0 ST Array (*42*) (GSE179417). DO mouse progressors in the study met endpoint morbidity criteria (most near 20 days post infection) whereas controllers remained above morbidity thresholds at 60 days post infection. In the third study DO mice were infected with 100 CFU M.tb HN878, and assigned as progressors or controllers based on a severity score including log_10_ CFU, area of lung inflammation and *a priori* threshold scores to segregate groups (*41*). In nearly all cases, all three risk signatures are able to significantly discern naïve animals from progressor groups (**Figure 2**). However, the differentiation between controllers and progressors was less robust across conditions or scores, and we hypothesize that this may be related to the infection dose. In the first Ahmed study, the NHP progressor group was infected with a ten-fold higher dose compared with the controller group, and samples were at a far later time point post infection for the progressors (22-24 vs 5-8 weeks). These differences likely influenced score performance heavily, since all scores were able to differentiate controllers from progressors, and controller scores were nearly comparable to naïve animals (**Figure 2 A-C**). All risk scores significantly correlated with lung bacterial burden and percent lung inflammation endpoints in this study (**Supplemental Figure 1**). Interestingly in both DO mouse studies, controller scores were much closer, albeit significantly different in some cases, to progressors than naïve animals and in all cases controllers displayed significantly elevated scores compared with naïve cohorts (**Figure 2 D-I**). In the case of DO mice in the Koyuncu *et al.* study, controllers displayed a significantly higher Sweeney3 score than progressors (**Figure 2 E**). Whereas in Ahmed *et* al, Risk6 and BATF2 were the discriminating signatures, differentiating controllers from progressor DO mice (**Figure 2 G, I**). There was a significant correlation between lung bacterial burden (CFU) and all three risk scores in the Ahmed DO mouse study (**Supplemental Figure 2 A, C, E**). Notably there was a lower correlation between risk scores and lung inflammation despite strong correlation between lung CFU and lung inflammation in the primary publication (**Supplemental Figure 2 B, D, F**). Correlations between area of lung inflammation and risk scores were weak (Risk6 R^2^ = 0.17, Sweeney3 R^2^ = 0.24, and Batf2 R^2^ = 0.15, graphs not shown), consistent with scores reflecting lung bacterial burden more than lung damage. Risk signatures derived from these transcriptional data sets begin to lose the ability to differentiate controllers versus progressors as the infection dose is increased (**Figure 2 D-I**), but scores still correlated with bacterial burden across samples (**Supplemental Figures 1-2**).

**Figure 2.**
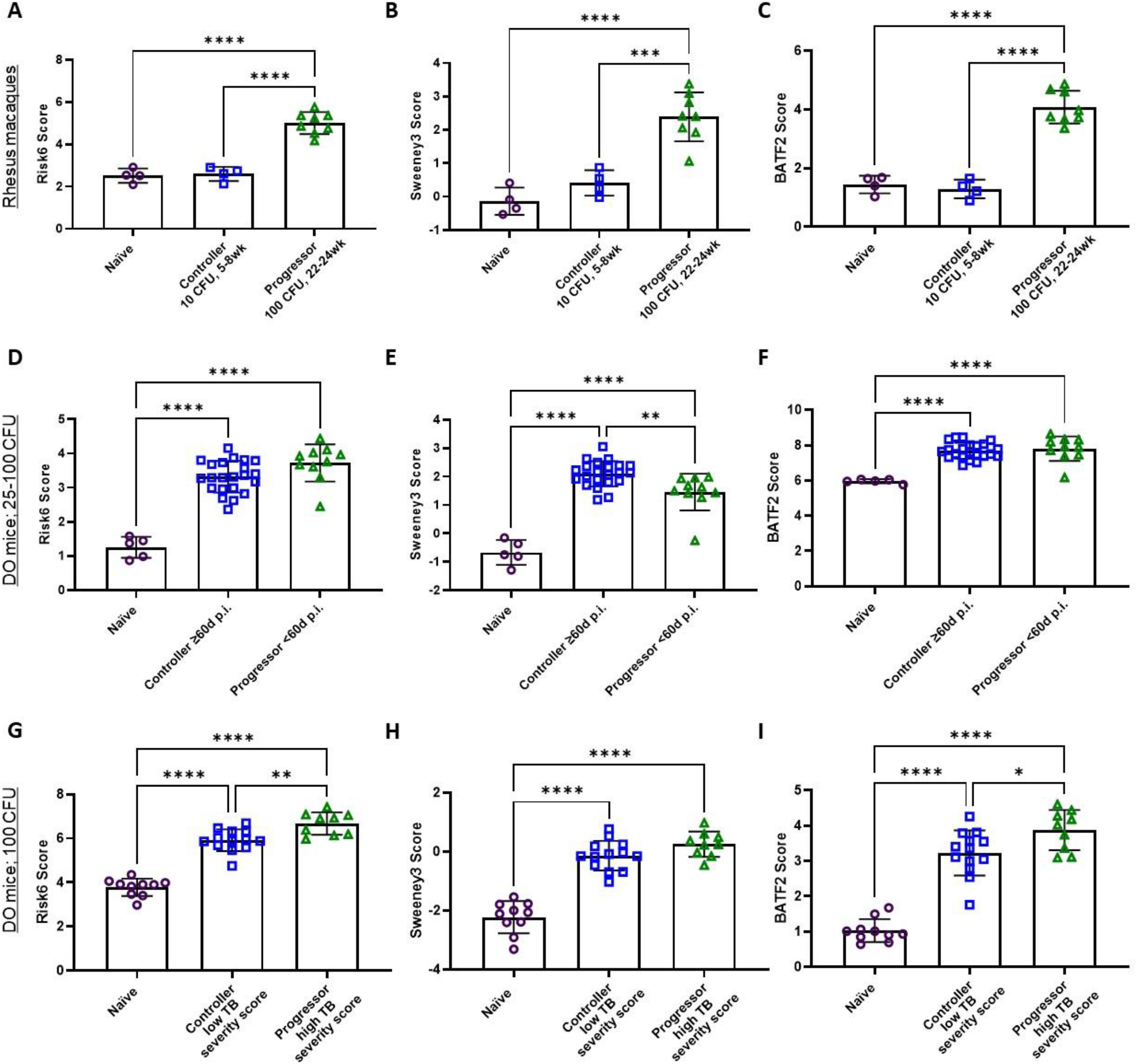
Risk signature scores assigned to naïve (open purple circles), controller (open blue squares) or progressor (open green triangles) NHP or DO mouse lung samples. **A-C**) From Ahmed *et al*, 2020, Indian rhesus macaques were infected with either 10 CFU and lung tissue obtained at 5-8 weeks post infection (controllers), or 100 CFU M.tb CDC1551 and tissue collected 22-24 weeks post infection (progressors) for RNA-seq. **D-F**) In Koyuncu *et al* (GSE179417) DO mice were infected with 25-100 CFU M.tb Erdman and lung RNA was analyzed by Affymetrix Mouse Gene 2.0 ST Array (GSE179417). Progressors met endpoint morbidity criteria and controllers persisted out to 60 days post infection. **G-I**) In Ahmed *et al,* 2020, DO mice were infected with 100 CFU M.tb HN878, and were assigned progressors or controllers based on “TB severity score” combining lung bacteria burden and inflammation. **A, D, G**) Risk6, **B, E, H**) Sweeney3 and **C, F, I**) BATF2 scores were calculated for each animal and cohorts were compared using one-way ANOVA with Tukey’s multiple comparison test correction. Significant comparisons are denoted by asterisks in the figure where * = p ≤ 0.05, ** = p ≤ 0.01, *** = p ≤ 0.001, **** = p ≤ 0.0001.

### A relatively high infection dose of M.tb drives high risk signature expression

In the final dose-related dataset, we explored whether there were identifiable differences in risk score performance in infection conditions over 100 CFU. Moreira-Teixeira and colleagues infected C57BL/6 and C3HeB/FeJ mice with “low” (100-450 CFU) or “high” (700-900 CFU) doses of M.tb HN878 or H37Rv (*43*). There were no significant differences between strain of M.tb or mouse (data not shown), therefore downstream analyses with these groups were aggregated into naïve, low- and high-dose cohorts. There was a consistently significant difference in the WB and lung tissue scores between naïve, and animals infected with the high or low dose of M.tb (**Figure 3**). Only Risk6 from the WB was able to differentiate between the low- and high-dose infection groups (**Figure 3 A**). The overlap observed here in distribution of risk scores may suggest that enough relatively higher (> 100 CFU) infections results in loss of resolution between groups when assessed at the same timepoints post infection.

**Figure 3.**
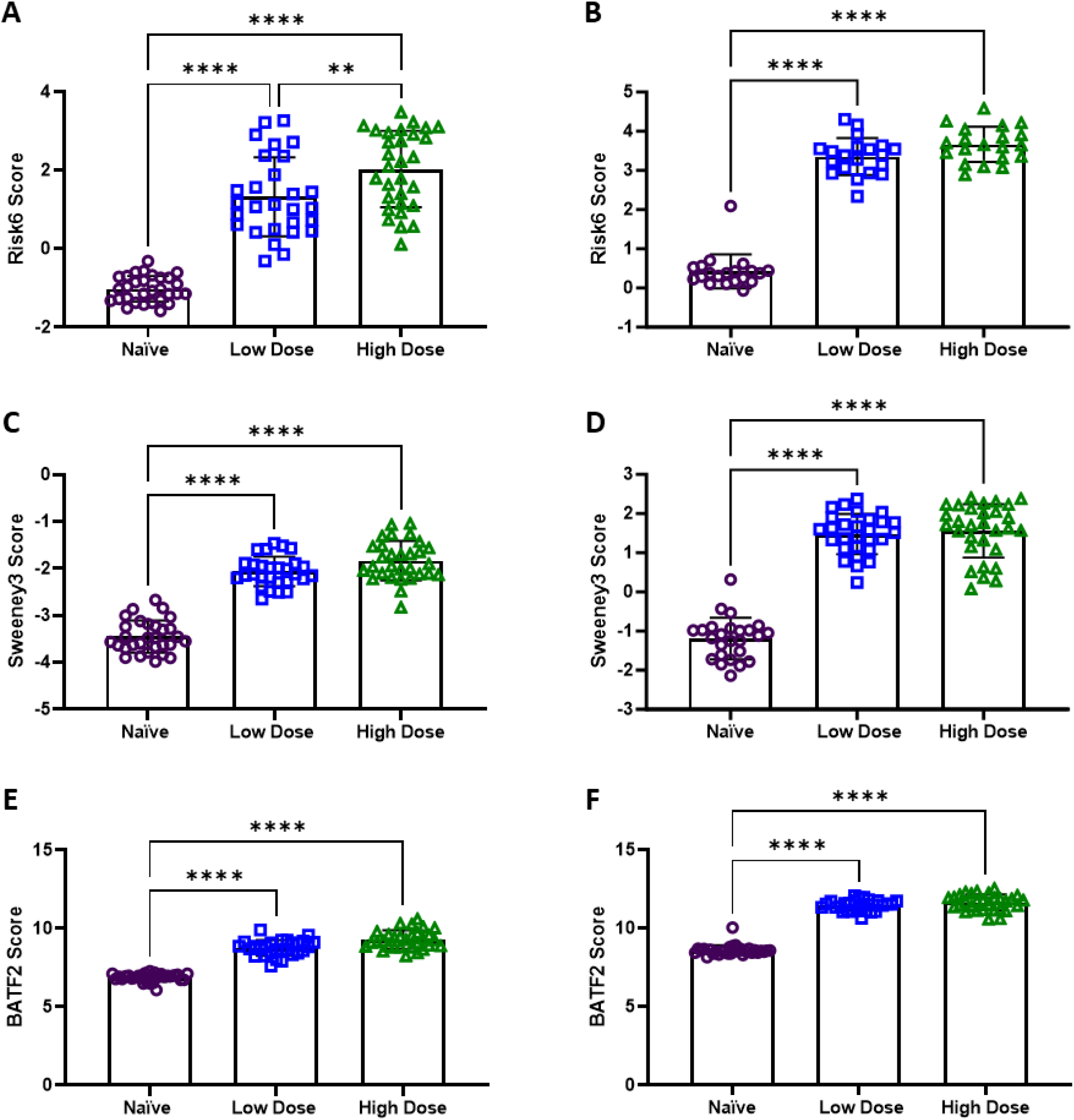
Risk signature scores assigned to naïve (purple circles), low dose (blue squares) or high dose (green triangles) C57BL/6 or C3HeB/FeJ mouse WB and lung samples. In Moreira-Teixeira *et al* C57BL/6 or C3HeB/FeJ mice were infected with a low dose (100-450 CFU) or high dose (700-900 CFU) of M.tb and **A, C, E**) WB (GSE137092) and **B, D, F**) lung samples (GSE137093) were collected for gene expression analysis by RNA-seq. **A, B**) Risk6, **C, D**) Sweeney3 and **E, F**) BATF2 scores were calculated for each animal and cohorts were compared using one-way ANOVA with Tukey’s multiple comparison test correction. Significant comparisons are denoted by asterisks in the figure where ** = p ≤ 0.01, **** = p ≤ 0.0001.

These data suggest that infection dose and progression of lung bacterial burden are the best predictors of risk score performance in WB and lung samples. Data available were largely collected between 3 and 22 weeks post infection, approaching or well within stationary/chronic stages of infection.

### Kinetics

We next investigated whether kinetics of infection would influence risk to active TB disease score performance. In humans, COR gene expression increases as an individual nears detectable disease progression, therefore we hypothesized we would observe similar kinetics in animal models as infections were established and bacterial burden increased over time. This reanalysis included ten datasets meeting our inclusion/exclusion criteria (**Table 4**). This kinetic category of reanalysis spans two species (mouse – BALB/c, C57Bl/6, C3HeB/FeJ, 129S2; and NHP – *Macaca fascicularis*), three tissue types (WB, lung and spleen), and involves five separate mycobacterial strains (*M. bovis*, M.tb H37Rv, M.tb Erdman and two Beijing isolates).

### Acute sampling post challenge reveals better correlation with risk signature expression over time

The kinetic data sets available included study designs with low- or high-dose infection with M.tb, the lowest being 25 CFU, and contained widely variable ranges of time. In order to examine broad changes or influences of time since infection or infection dose, the Pearson correlation R^2^ value for each data set (correlation between risk signature and day post infection) was calculated and graphed versus median days post infection (**Figure 4, Supplemental Table 3**). We did not observe trends in correlations based on infection dose (**Figure 4**). However, it does appear that R^2^ values were higher in data sets with lower median days post infection, particularly for Risk6 (**Figure 4 A**) and Batf2 (**Figure 4 C**) signatures. Six datasets included a baseline Day 0 sampling and when Day 0 average scores were used to normalize subsequent day scores, there was no further resolutions of correlation between score and time since infection (data not shown).

**Figure 4.**
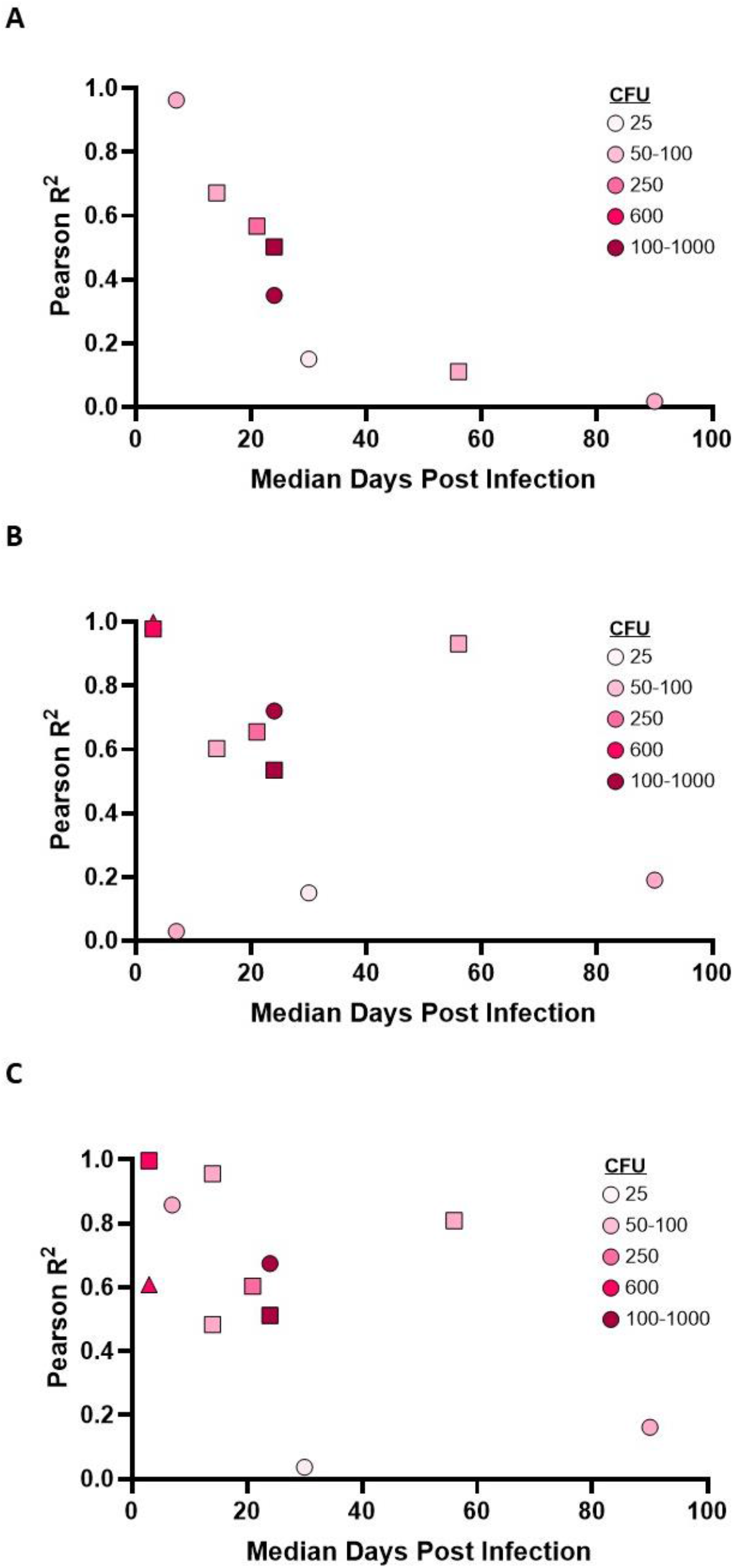
Pearson correlation R^2^ of risk signature scores compared with time since infection. **A**) Risk6, **B**) Sweeney3 and **C**) BATF2 scores were calculated for each animal and a Pearson correlation was run evaluating the signature score versus time since challenge. Challenge dose is denoted by the color intensity of the shape (CFU 25 low to 100-1000 high). Samples derived from blood are denoted as circles, samples from lung are denoted as squares and those derived from spleen are triangles. Accompanying p values are listed in Supplemental Table 3.

### Samples derived from lung in preclinical studies is an option for specific signatures

Moreira-Teixeira and colleagues’ study design includes BALB/c mice that were infected with 100-1000 CFU M.tb H37Rv and contain both lung- (GSE140944) and blood-derived (GSE14943) data over time. RISK6, Sweeney3 and BATF2 scores were used to examine whether blood- and lung-derived signatures performed similarly or disparately over time (**Figure 5**). Overall, lung-derived tissue scores (**Figure 5 B, D, F**) showed a greater increase over time than matched signature blood-derived scores (**Figure 5 A, C, E**). We also noted that Sweeney3 scores derived from blood did not increase over time and performed poorly compared with other scores (**Figure 5 C**). For Risk6 and Batf2 signatures the lung- and blood-derived scores started to differentiate from baseline around 28-days post infection (**Figure 5 A, B, E, F**). These data suggest that both blood, the traditional human sampling strategy, and lung, could be viable tissue types for evaluating COR signatures in preclinical models of TB disease. Furthermore, given the lung is the primary site of infection and disease it is not surprising that the resolution of scores from this tissue may outperform blood-based signatures in some conditions.

**Figure 5.**
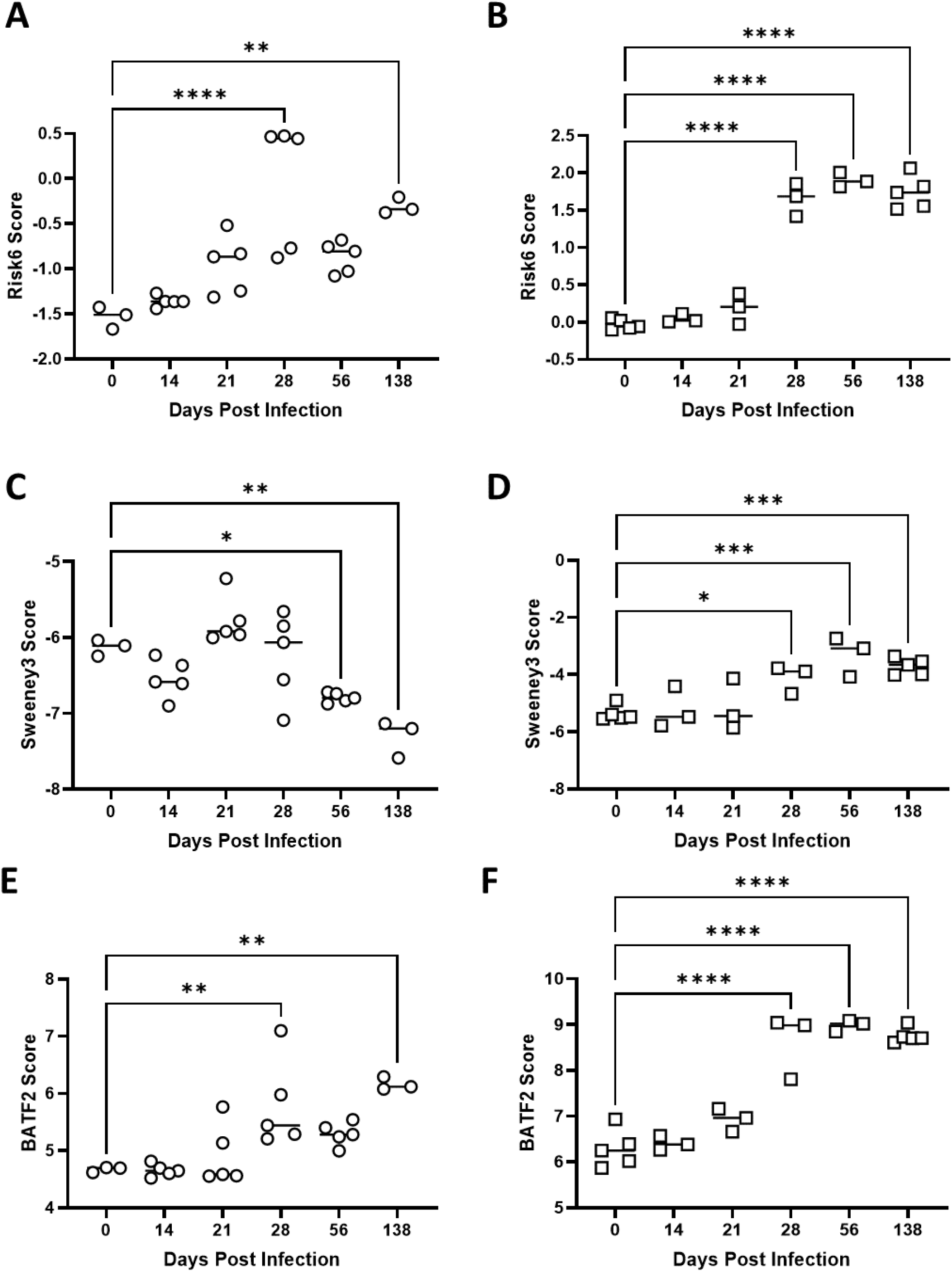
Risk signature scores derived from blood (open circles, **A, C, E**) or lung (open squares, **B, D, F**) tissue samples. In Moreira-Teixeira *et al* (GSE140943 blood, left and GSE140944 lung, right) BALB/c mice were infected with a high dose (100-1000 CFU) of M.tb H37Rv. Samples were evaluated for RNA analysis using the Illumina MouseWG-6 v2.0 expression beadchip and **A, B**) Risk6, **C, D**) Sweeney3 and **E, F**) BATF2 scores were calculated for each animal. Scores were compared using one-way ANOVA with Dunnett’s multiple comparison test correction to tissue matched day 0 scores. Significant comparisons are denoted by asterisks in the figure where * = p ≤ 0.05, ** = p ≤ 0.01, *** = p ≤ 0.001, **** = p ≤ 0.0001.

### Kinetic evaluations of score performance requires high resolution between groups

The final dataset recovered from our search was by Gideon *et al*., where Cynomolgus macaques were infected with an instillation of 25 CFU M.tb Erdman and followed for up to 180 days post infection. WB was collected for RNA and analyzed by an Illumina HumanHT-12 V4.0 expression beadchip (*49*). NHPs were differentiated into active or latent TB groups at the termination of the study and retrospectively analyzed based on clinical diagnosis and intensity of pulmonary lesions with 18F-fludeoxyglucose (FDG) and imaging. Using calculated risk signature scores, we observed that only the active disease group had a weak correlation (Spearman r = 0.5934, p = 0.0360) with time since infection, but no significant difference within the active or latent groups over time (**Figure 6 A**). Similar to the authors of the original study, we noted that the most robust increase in Risk6 score was observed day 20 through day 56 post infection compared with day 0. Surprisingly, there was no correlation between risk scores and FDG avidity scores. Together these data suggest that the risk scores, in this experiment, were not strongly associated with disease severity, nor are they able to differentiate active versus latent categorically by these endpoints.

**Figure 6.**
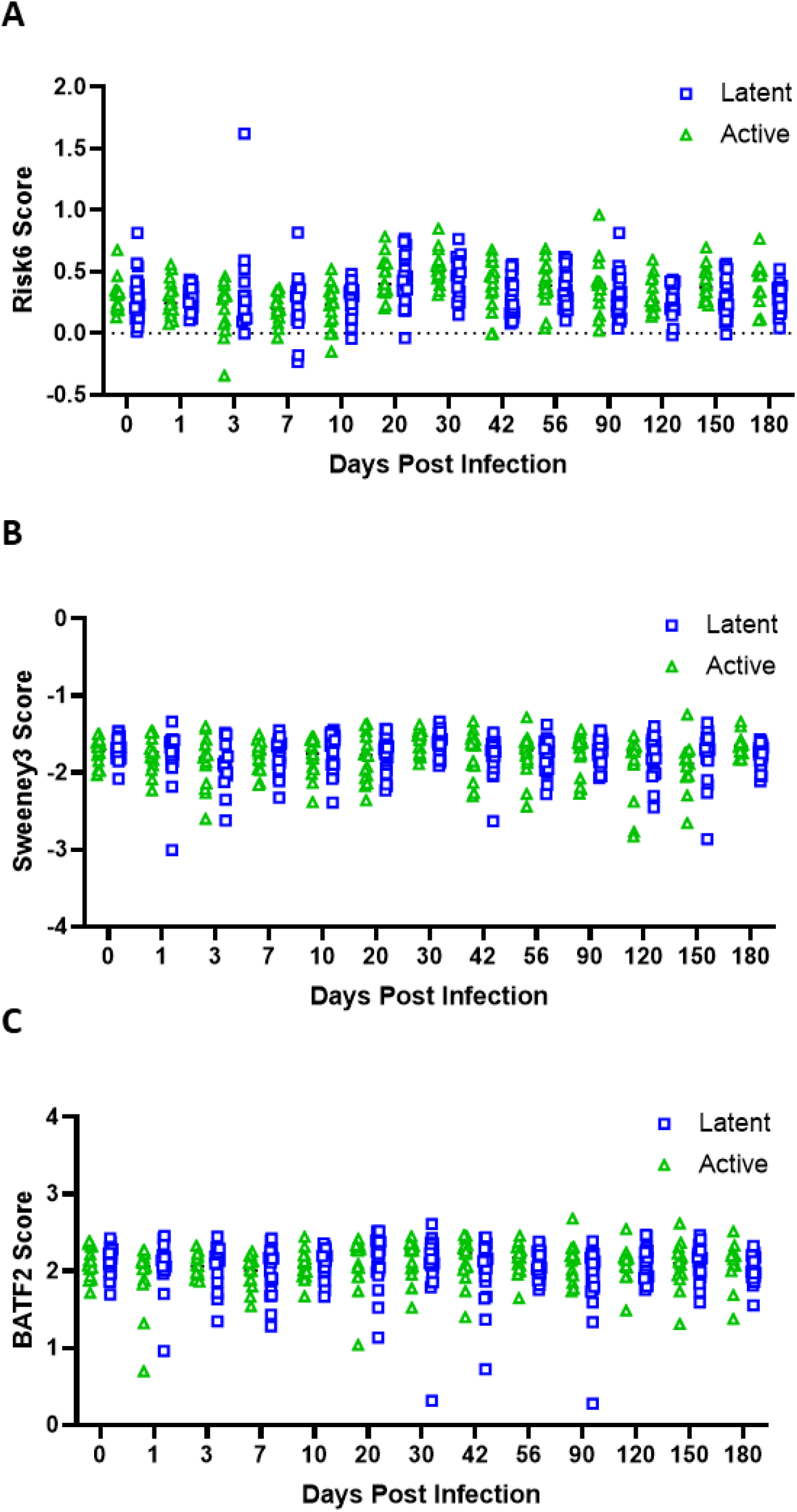
Risk signature scores derived from WB samples from Cynomolgus macaques with either latent (open blue squares) or active TB disease (open green triangles). In Gideon *et al* (GSE84152) animals were infected with an instillation of 25 CFU M.tb Erdman and samples were taken from pre-infection through day 180 post-infection. Gene expression analysis was performed by Illumina HumanHT-12 V4.0 expression beadchip. **A**) Risk6, **B**) Sweeney3 and **C**) BATF2 scores were calculated for each animal and separate comparisons of active and latent cohorts were performed using one-way ANOVA with Tukey’s multiple comparison test correction.

In line with the dose data, these data suggest that infection dose and progression of lung bacterial burden are the best predictors of risk score performance in the WB and lung samples and too high of an initial infection reduces the predictive ability of the risk signatures evaluated here. However, lung-derived samples appear more predictive than blood-based samples in some cases, and should be followed up in future work.

### Interventions

Most therapeutic treatment is designed to reduce *in vivo* bacterial burden either through direct bactericidal activity or host-mediated control. Therefore, we hypothesized that we would observe decreasing COR gene expression through courses of successful drug treatment or vaccination. Conversely, genetic interventions or treatment which resulted in expanding bacterial burden or dampened host-immunity would subsequently result in higher COR scores. This reanalysis included 14 datasets meeting our inclusion/exclusion criteria (**Table 5**). This intervention category of reanalysis spans two species (mouse – C57Bl/6, Sp110, Sp140, A/Sn, I/St, ΔdblGata, DBA/2, BALB/c, B6D2F1, C3HeB/FeJ; and NHP – *Macaca mulatta*), three tissue types (blood, lung and spleen), and involves four M.tb strains (Erdman, H37Rv, HN878 and CDC1551).

### Some mechanistic (genetic, depletion or metabolic) interventions influence signature gene expression

Deletion or depletion of key immune regulators are known to enhance preclinical model susceptibility to M.tb infection and allow higher bacterial burden. Interestingly, assessment of risk signature expression from lung samples derived from TB-susceptible I/St or TB-resistant A/Sn mice (*66*) in a study by Wilk and colleagues (GSE64167) did not reveal significant differences (data not shown). Similarly, in Bohrer *et al* (GSE165871) mice lacking eosinophils, ΔdblGata, demonstrated equal Risk6, Sweeney3 and BATF2 scores from lung RNA samples compared to C57BL/6 mice after challenge with M.tb H37Rv (data not shown). The interest in connecting genetic loci to disease outcomes in mice has also revealed the super susceptibility to tuberculosis 1 (Sst1) locus. In the study by Ji and colleagues (GSE166114) there are trends (n=2) of higher Risk6, Sweeney3 and BATF2 signature expression in RNA samples from M.tb Erdman challenged Sp140^-/-^ mice (lacking a repressor of type I IFN transcription) compared to C57BL/6 mice (data not shown). In the same study the reverse trend is observed for Sp110^-/-^ mice (lacking NF-κβ modulating nuclear protein), where all signatures are lower than C57BL/6 mice (data not shown). Studies evaluating downstream immune effectors may further help identify the relationship between key immune regulators and risk signature expression during M.tb infection. For example, granulocyte-macrophage colony-stimulating factor (GM-CSF) has been shown to be an influential cytokine involved in the inhibition of intracellular M.tb growth (*67*). Moreira-Teixeira and colleagues (GSE141207) collected RNA from blood and lung tissue from C57BL/6 mice given either α-GM-CSF or isotype control, and subsequently infected with 200 CFU M.tb HN878. Infected cohorts, regardless of treatment group, harbored significantly higher risk signatures than naïve mice in both tissues (data not shown). When comparing risk signatures from infected animals we observed that all three signatures were more affected by α-GM-CSF treatment in blood samples (**Figure 7 A-C**) than in samples derived from lung tissue (**Figure 7 D-F**). In lung samples, there was no difference observed in the Sweeney3 signature (**Figure 7 E**), whereas all other conditions demonstrated a significant increase in score with α-GM-CSF treatment (**Figure 7**). This difference between tissues and changes in risk signature expression may be partially due to penetrance of α-GM-CSF antibodies in solid tissue versus peripheral blood and the magnitude of changes that may subsequently occur. Tumor necrosis factor (TNF) is another important cytokine involved in the control of M.tb *in vivo* (*68–70*). In a study by Dutta *et al*. (GSE55183), they compared transcriptional differences from C3HeB/FeJ mice infected with M.tb H37Rv with or without α-TNF treatment. In lung samples derived from these experiments, *in vivo* depletion of TNF shows a trend of increasing COR signature expression, statistics were not performed since the group size (n=2) for this series was too small (**Figure 7 G-I**). These data suggest that in the same way reducing key cytokines can enhance susceptibility to M.tb in humans with immunosuppressive therapy, preclinical *in vivo* depletion of influential cytokines can increase TB disease risk signatures.

**Figure 7.**
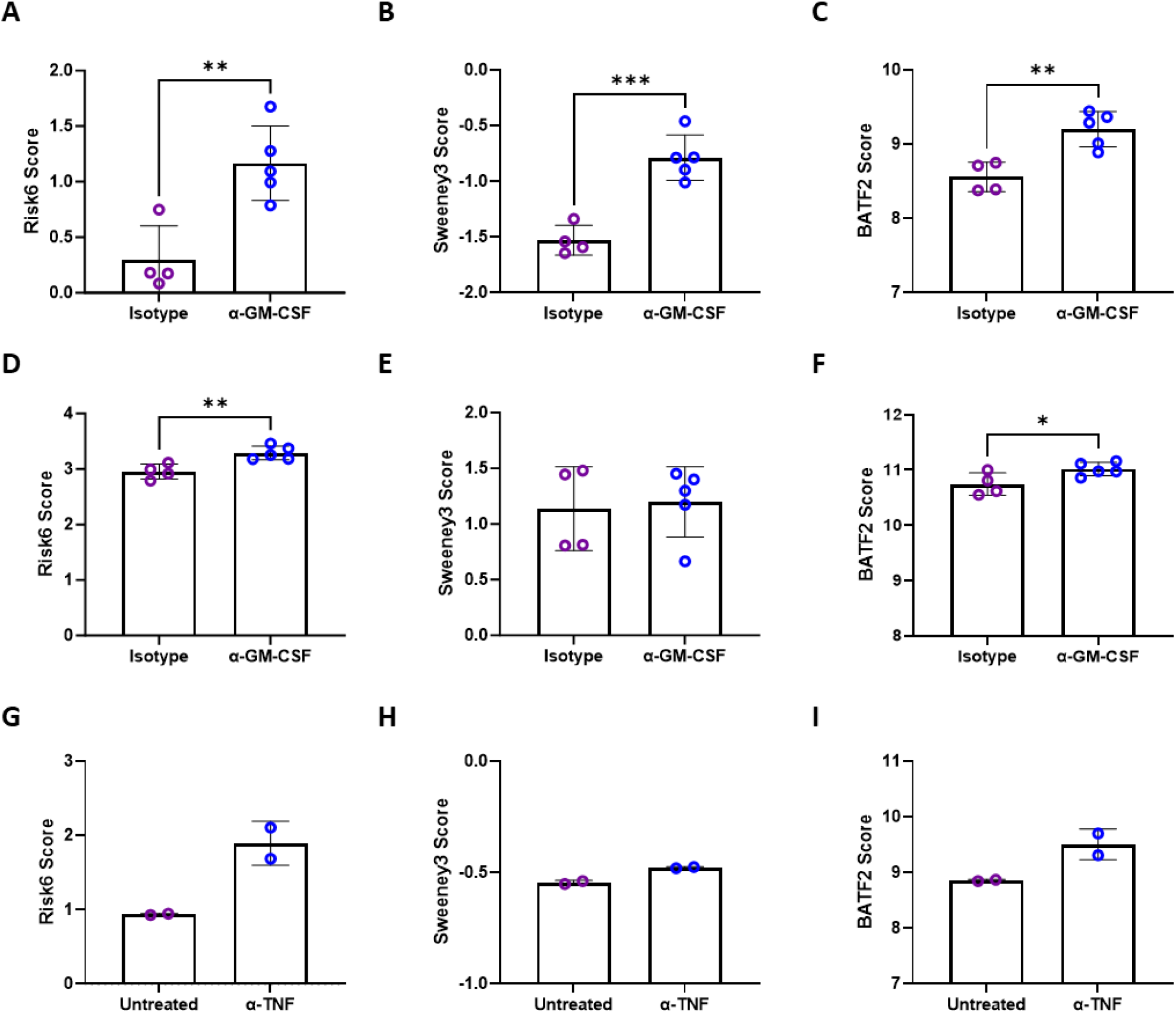
Risk signature scores increase with depletion of cytokines involved in M.tb control. **A-F**) In Moreira-Teixeira *et al* (GSE141207) C57BL/6 mice infected with 200 CFU M.tb HN878 and treated with either an isotype control (open purple circles) or α-GM-CSF depleting antibody (open blue circles) one day before infection and twice per week during the study. RNA from both **A-C**) blood and **D-F**) lung were collected for sequencing (GSE141207). **G-I**) In Dutta *et al* (GSE55183) C3HeB/FeJ mice were infected with ∼1.0 OD_600_ M.tb H37Rv. After six weeks mice were left untreated (open purple circles) or given an α-TNF depleting antibody (open blue circles) twice per week for four weeks and lung tissue was collected for RNA analysis (GSE55183). **A, D, G)** Risk6, **B, E, H**) Sweeney3 and **C, F, I**) BATF2 scores were calculated for each animal. In **A-F** groups were compared by t-test, but in **G-I** groups were not statistically compared due to low n of 2 per group. Significant comparisons are denoted by asterisks in the figure where * = p ≤ 0.05, ** = p ≤ 0.01 and *** = p ≤ 0.001.

Given the significant roles of immunometabolism on host-pathogen outcomes we hypothesized that metabolic interferences may exacerbate M.tb infection and likewise influence genes in the POD risk signatures being evaluated. These data are particularly of interest given the recent success of the RATIONS clinical trial in Jharkhand India (reducing Activation of Tuberculosis by Improvement of Nutritional Status; Clinical Trial Registry of India: CTRI/2019/08/020490), where nutritional support to households with a member with active TB reduced incidence and subsequent infections significantly (*71*). This series of reanalysis includes RNA samples derived from spleen and lung tissues of mice infected with M.tb undergoing metformin treatment, caloric restriction or treatment intended to lower triglycerides. Metformin given for 30 days reduced Risk6 and BATF2 signature scores compared with similarly infected but untreated mice (**Figure 8 A, C**) in data from Singal and colleagues (GSE57275). This trend was not observed for Sweeney3 (**Figure 8 B**). Conversely, in data from Matarese *et al.* (GSE127263), caloric restriction increased Sweeney3 scores when compared with chow given *ad libitum* in mice infected with M.tb, and not the other two scores tested, despite caloric restriction having a protective effect against bacterial burden *in vivo* as reported by the authors (**Figure 8 D-F**). Gandotra *et al.* (GSE179437) treated M.tb infected C3HeB/FeJ mice from day 8 to 28 post infection with compound T863, designed to inhibit triglyceride synthesizing enzyme DGAT1. In RNA derived from the lungs of these infected mice treatment with T863 did not change signature score expression compared with untreated mice, even when segregated by low (100) or high (500 CFU) infection (**Figure 8 G-I**).

**Figure 8.**
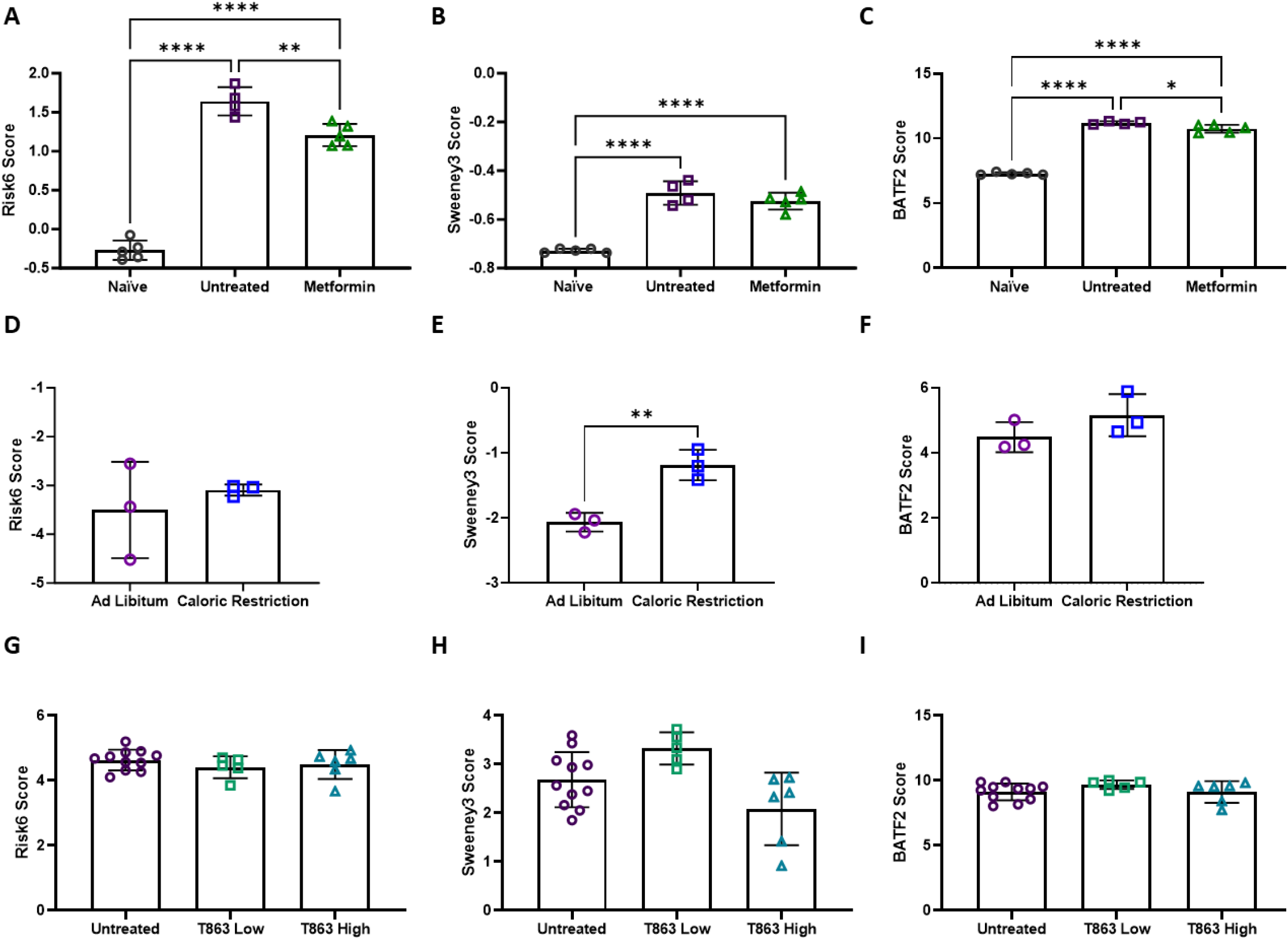
Risk signature scores are moderately influenced by metabolic interventions. **A-C**) In Singal *et al* (GSE57275) C57BL/6 mice were infected with 10^3^ CFU M.tb H37Rv for 7 days and left untreated (open purple squares) or treated with metformin (250mg/kg, open green triangles) six days a week for 28 days and lung tissue from these cohorts and naïve cohorts (open black circles) was collected for RNA analysis using Illumina MouseWG-6 v2.0 expression beadchip (GSE57275). **D-F**) Matarese and colleagues (GSE127263) infected DBA/2 mice with 10^5^ CFU M.tb H37Rv by i.v. These animals were given *ad libitum* normal chow (open purple circles) or chow intended to result in caloric restriction (open blue squares). Diets were maintained for 40 days up until spleen tissue was collected for RNA-seq analysis. **G-I**) In Gandotra *et al* (GSE179437) C3HeB/FeJ mice were infected with 100 (low, open green squares) or 500 (high, open teal triangles) CFU M.tb Erdman. T863 was administered every three days from day 7 up until 28 via i.v. when lung tissues were collected and used for RNA-seq analysis along with untreated (open purple circles) animals (GSE179437). **A, D, G)** Risk6, **B, E, H**) Sweeney3 and **C, F, I**) BATF2 scores were calculated for each animal and cohorts were compared using an unpaired t-test where two groups are compared or an ordinary one-way ANOVA with Tukey’s multiple comparisons test where three or more groups are compared. Significant comparisons are denoted by asterisks in the figure where * = p ≤ 0.05, ** = p ≤ 0.01, **** = p ≤ 0.0001.

### TB therapy reduces COR scores in small animal models

This series of reanalysis includes RNA samples derived from lung tissues of mice infected with M.tb only and is an interesting examination of the site of infection and score performance. In highly effective and frontline antibiotic studies, decreasing Risk6 and BATF2 signatures correlate with treatment (**Figure 9**). In the case of dual therapy with rifampicin (R) and isoniazid (H) for 30 days during the chronic phase of infection (> 30 days post infection), Rodrigues and colleagues (GSE44825) observed no detectable CFU after this treatment. Reanalysis of gene expression data from this study demonstrated that Risk6 and BATF2 scores are significantly decreased in treated cohort versus the untreated cohort (**Figure 9 A-B**). Similarly, four weeks of RHZE (rifampicin, isoniazid, pyrazinamide and ethambutol) treatment lowered both Risk6 and BATF2 scores but not Sweeney3 in a study by Dutta *et al*. (**Figure 9 C-E**). Pyrazinamide administered as a monotherapy for 35 days beginning at 28 days post infection reduces all three signatures compared with untreated animals (**Figure 9 F-H**).

**Figure 9.**
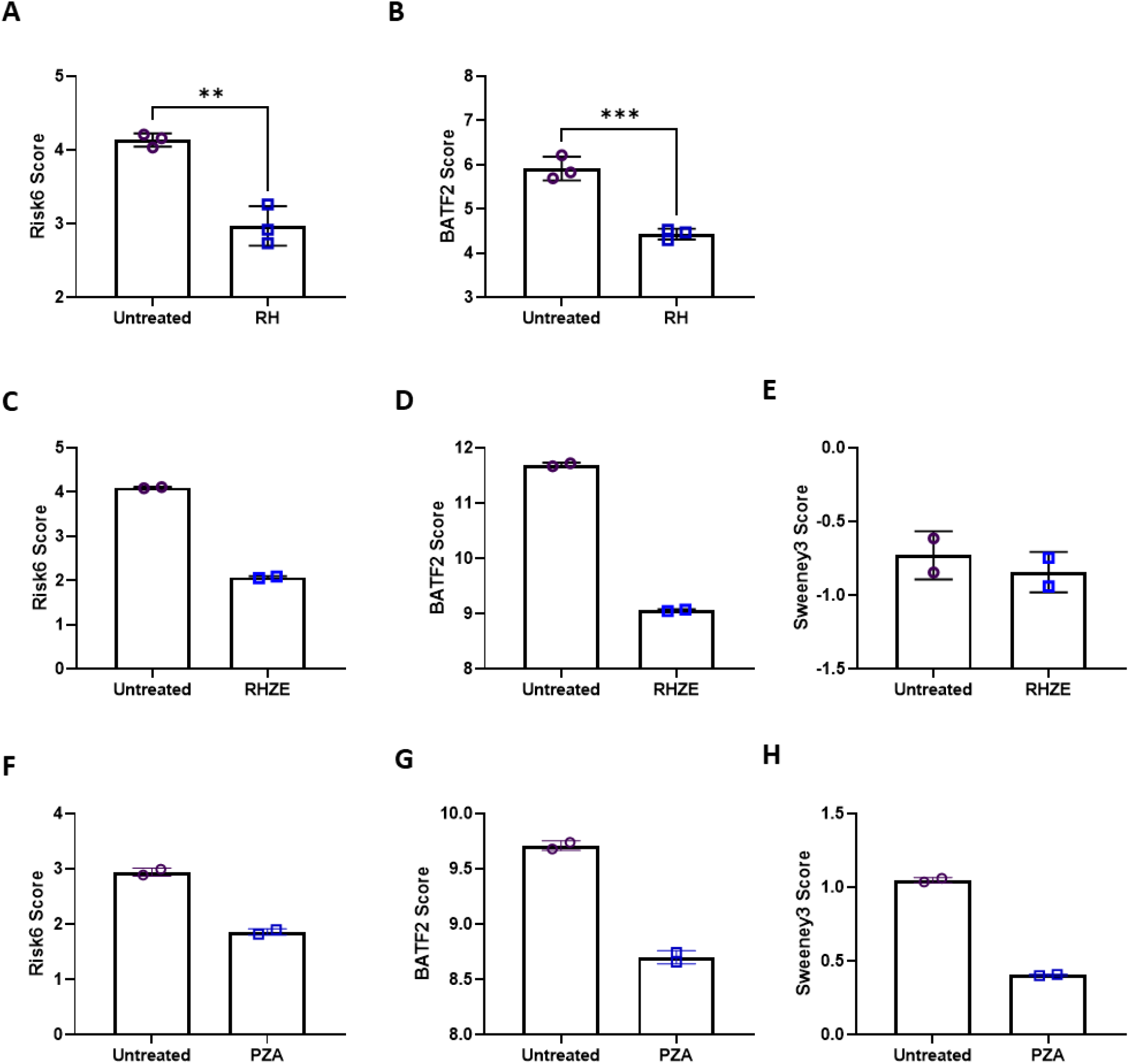
Risk signature scores from expression data derived from lungs of untreated (open purple circles) or drug treated (open blue squares) mice. **A-B**) In Rodrigues *et al* (GSE44825) BALB/c mice infected with 10^5^ CFU M.tb H37Rv for 30 days and left untreated or treated for 30 days with rifampicin 20 mg/kg and isoniazid 25 mg/kg (RH) for 30 days. **C-E**) Dutta and colleagues (GSE99625) infected BALB/c mice with 5x10^3^ CFU M.tb H37Rv and six weeks after infection treated with rifampicin 10 mg/kg + isoniazid 10 mg/kg + pyrazinamide 150 mg/kg + ethambutol 100 mg/kg (RHZE) orally for 4 weeks. **F-H**) In a study by Manca *et al* (GSE48027) B6D2F1 mice were infected with 250 CFU M.tb CDC1551 then given pyrazinamide 150 mg/kg (PZA) for 35 days. **A, C, F)** Risk6, **B, D, G**) BATF2 and **E, H**) Sweeney3 scores were calculated for each animal and cohorts were compared using an unpaired t test for **A-B** only where n=3. Significant comparisons are denoted by asterisks in the figure where * = p ≤ 0.05, ** = p ≤ 0.01, *** = p ≤ 0.001. Scores were calculated for each animal, but not statistically evaluated when n = 2 per group (**C-H**).

The above treatment regimens (RH, RHZE and PZA) are known to reduce pulmonary bacterial burden and the risk signatures appear to track well with this trend. This holds true when treatments are either non-effective or sub-optimal and result in ‘relapse-like’ chronic infections. For example, Subbian *et al*. investigated whether compound CC-11050, a phosphodiesterase-4 inhibitor, would synergize with isoniazid (H) treatment (*61*). Importantly, CC-11050 treatment alone does not reduce bacterial burden compared with untreated mice in published literature and similarly there is no difference in risk signature scores between these cohorts. However, there is an expected and significant increase in signature scores between naïve samples and both infection cohorts (**Figure 10 A-C**). We observe in Park *et al*. (GSE97835) that end of treatment (EOT) with HZE timepoints with persistent bacteria retain higher signature scores than naïve samples. After alleviation of treatment, wherein bacteria are expected to expand again, signature scores increase (post EOT) (**Figure 10 D-F**).

**Figure 10.**
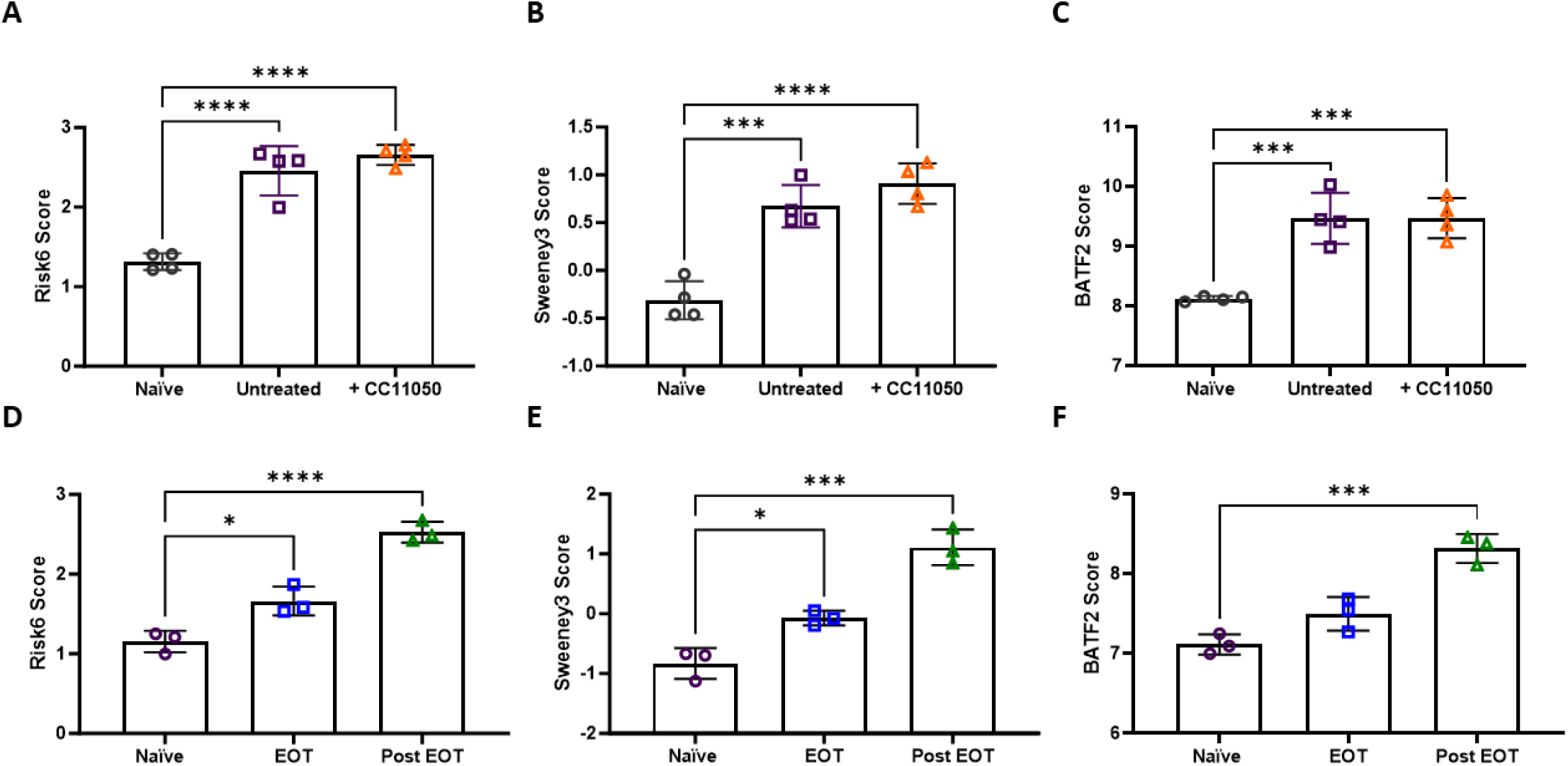
A-C) Risk signature scores calculated from naïve (open black circles), infected but untreated (open purple squares) and infected and CC11050 treated (open orange triangles) B6D2F1 lung samples. In Subbian *et al* B6D2F1 mice were infected with 100-150 CFU M.tb CDC1551 and lung samples were collected for microarray analysis (GSE83188). **D-F**) Risk signature scores calculated from naïve (open purple circles), end of treatment (EOT, open blue squares) or post treatment (Post EOT, open green triangles) C57BL/6 lung samples. In Park *et al* C57BL/6 mice were infected with 200-300 CFU M.tb Erdman and lung samples were collected for microarray analysis. Samples were collected after treatment with a daily dose of INH: 26.5 ± 0.9 mg/kg; PZA and EMB: 132.6 ± 4.7 mg/kg at four weeks after treatment cessation (GSE97835). **A, D**) Risk6, **B, E**) Sweeney3 and **C, F**) BATF2 scores were calculated for each animal and cohorts were compared using one-way ANOVA with Tukey’s multiple comparison test correction. Significant comparisons are denoted by asterisks in the figure where * = p ≤ 0.05, *** = p ≤ 0.001, and **** = p ≤ 0.0001.

### BCG vaccination in NHP demonstrates a protective effect against COR signature score expression

The final category of intervention-based datasets available included data from Hansen and colleagues (*65*) where rhesus macaques were vaccinated with BCG, RhCMV/TB candidate or a prime-boost strategy of BCG with RhCMV/TB. RNA was extracted from WB in Paxgene tubes and analyzed by RNA-seq. After vaccination, NHPs were infected with 10-25 CFUs of M.tb Erdman. In all groups, the Risk6 scores increased significantly by 28 days post infection compared with day 0 (day 0 v 28 untreated p < 0.0001; RhCMV/TB p = 0.0006; BCG p = 0.0136; BCG + RhCMV/TB p = 0.0041). Additionally, when evaluating within day 28, groups with BCG regimens displayed slightly lower, but still significant, signatures compared with the untreated cohort at the same timepoint (**Figure 11 A**). Similarly, Batf2 scores were significantly higher for all of the groups by day 28 when compared with both day 0 and day 14 post infection (day 0 v 28 Untreated p < 0.0001; RhCMV/TB p < 0.0001; BCG p = 0.0001; BCG + RhCMV/TB p = 0.0006). For Batf2 scores at day 28, groups with BCG regimens displayed significantly lower signatures compared with the untreated group (**Figure 11 B**). Each of the NHPs also had corresponding harmonized necropsy disease scores which were made up of the pathologic disease observed and recovered M.tb from tissue. These necropsy scores did not correlate significantly with COR signature score (neither by group nor aggregated) (**Figure 11 C-D**).

**Figure 11.**
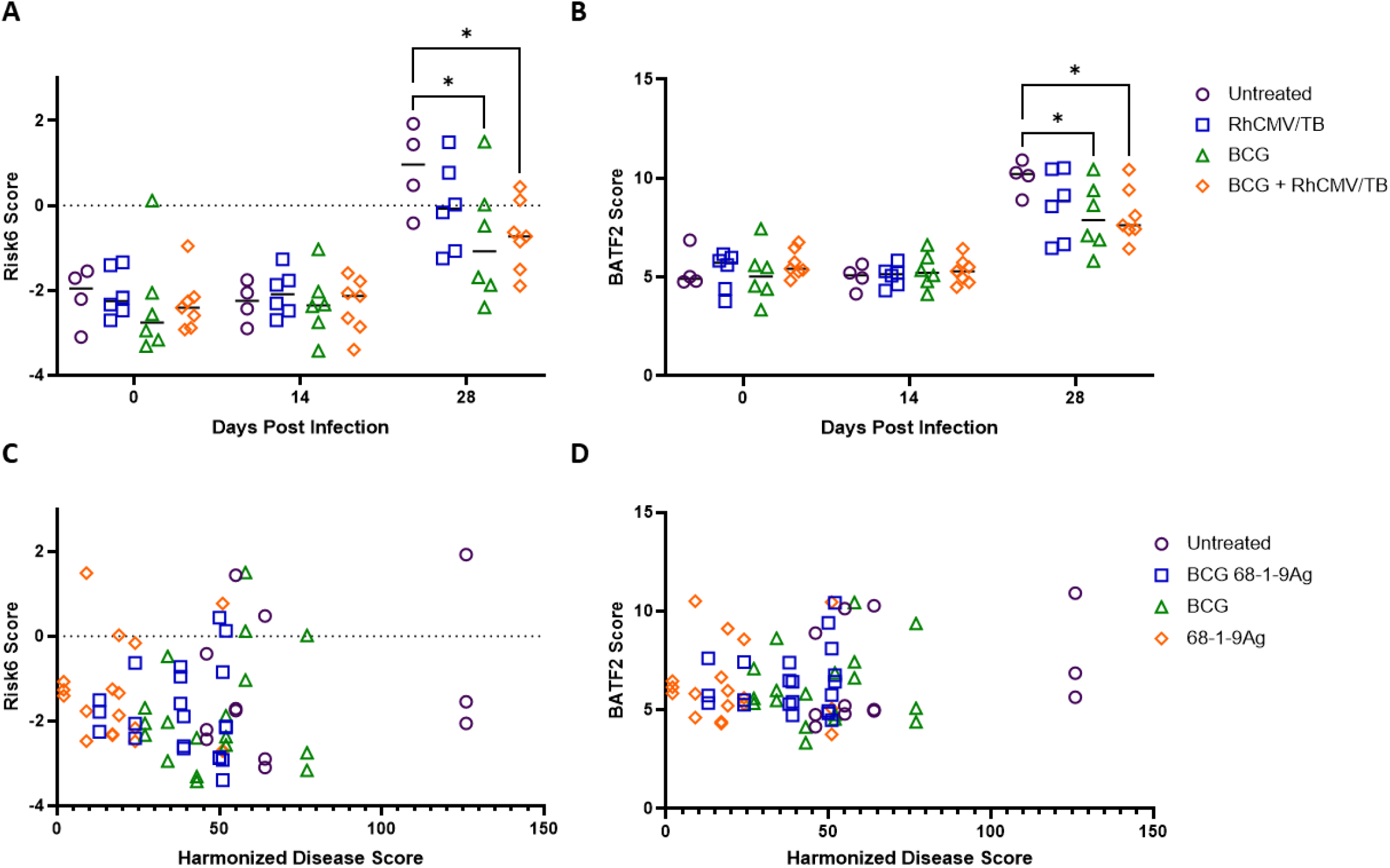
Risk signature scores derived from WB at 0, 14 or 28 days post infection with M.tb Erdman. In Hansen *et al* (GSE102440) cohorts of Rhesus macaques were left untreated (open purple circles), or vaccinated with BCG 68-1-9Ag (open blue squares), BCG (open green triangles) or 68-1-9Ag (open orange diamonds) and subsequently infected with 10-25 CFU M.tb Erdman and followed over time. **A**) Risk6 and **B**) BATF2 scores were calculated for each animal and cohorts were compared at each timepoint using a two-way ANOVA with Tukey’s multiple comparison test correction. Significant comparisons are denoted by asterisks in the figure where * = p ≤ 0.05. **C**) Risk6 and **D**) BATF2 scores were evaluated with harmonized disease score for correlations using a Spearman-rank correlation test.

These intervention-based data show particularly a strong association between vaccine or drug treatment outcomes with COR gene signature scores. This may imply that these transcriptional endpoint tools could be used for early stage-gating of novel therapies against tuberculosis.

## DISCUSSION

In this reanalysis of public datasets we hypothesized that human-derived risk signature scores would be able to segregate naïve from infected animals, and stratify along with bacterial burden and/or disease burden endpoints in preclinical models of M.tb infection. The relative heterogeneity of immune responses and TB disease progression in different human clinical demographic groups (including unknown time since infection, when or how swift individuals will progress to active disease) still has continued to result in numerous predictive or diagnostic gene signatures. Given animal studies have less intrinsic variability and known infection time and dose, we felt this investigation would be low-risk high-reward for the field. These data collectively support that Risk6, Sweeney3 and Batf2 risk signatures can be used as transcriptional endpoints in preclinical animals and that resolution is improved in specific conditions. Divergence between study designs, including dose and kinetic timepoints of sampling at first seemed a limitation of this reanalysis, but has instead provided novel hypotheses about traditional endpoints that best correlate with signature scores. For example, lower infection doses invoke lower signature expression over time and can induce more variability across the group in line with trends in bacterial burden in these cohorts. In general, bacterial burden was found to more consistently correlate with higher signature gene expression compared with pulmonary immunopathology, imaging or overt disease scoring. Kinetic data sets suggest that RNA samples derived from lungs had better performance than WB samples overall. Additionally, early timepoints offered higher resolution than chronic timepoints captured, which is in line with lower initial bacterial deposition leading to a greater heterogeneity and resolution of scores. Conversely, we hypothesize that with a higher initial infection dose the COR signature kinetics are disrupted or reaching an upper limit, making discrimination between groups more difficult. The mechanistic, drug and vaccine-based intervention results support using COR signatures as endpoints for future evaluation of therapeutic candidates in the preclinical pipeline.

One major limitation of these findings is that none of the studies in which the datasets were obtained from for reanalysis were originally designed to address hypotheses related to COR score performance or identify confounding influences of study designs on COR scores. Follow on studies designed and powered to do so are warranted given the proof-of-concept results presented here. For example, these studies individually or aggregated are not powered to evaluate the contribution of the M.tb strain or lineage on signature gene expression or specific correlations with other endpoints. This remains an interesting question in both clinical and preclinical settings. Highly virulent M.tb strains may promote enhanced COR gene expression given their relatively advanced kinetics of *in vivo* expansion. Sequencing of microbiologically confirmed human TB cases within clinical studies could also help address this question, however sequencing is not routinely performed in many resource limited areas at present. A second limitation of this analysis is that while data evaluated here was comprised of both male and female animals in some studies no sets of metadata included which data were derived from either sex. Given recent and exciting interest in sex-derived differences in TB disease and immune responses (*72–74*), evaluating whether gene signatures also diverge based on sex is worthwhile.

Future and active work in COR signatures are interrogating the inherent mechanisms and networks of gene pathways to further understand the implications of score performance. Interestingly, Gupta and colleagues have previously used Ingenuity Pathway Analysis to design a network of upstream gene or cytokine regulators of several COR gene signatures, including the three evaluated here (*32*). In those analyses, a number of genes in RISK6, Sweeney3 and BATF2 feature prominently in *STAT1* and *IFNG* pathways, but not all. We hypothesize that the divergences are what differentiate score performance within sample sets as observed here. As more work is done to determine whether COR genes are directly tied to the mechanisms of disease progression or are merely an associated readout, having homologous signatures may accelerate the cross-pipeline validations and discoveries. Since these scores have shown promise in samples derived from *in vivo* preclinical models, originally excluded single-cell RNA-seq data could be leveraged to evaluate which cells most contribute to signature gene expression across expansive *in vitro* and *ex vivo* conditions, generating and validating novel hypotheses. The data evaluated here was exclusively derived from mouse or NHP-derived tissues. Several rabbit data sets were identified in our searches, but true orthologs of selected human gene sets of interest are difficult to confirm with confidence. Intermediate models such as guinea pigs and rabbits which display more similarities of human disease pathology than mice, but are less expensive and more readily available that NHPs, need more immunologic tools to enter the field as it exists presently. However, once these needs are met, we expect that both guinea pigs and rabbits will become valuable validation benchmarks for COR signatures as stage-gating endpoints. We predict that novel signatures will be identified that are high performing but unique to specific species due to innate model differences or non-overlapping gene orthologs. These novel species-specific biomarkers are highly valuable for the preclinical TB vaccine pipeline and should be pursued, but may be limited in their inherent translation. These data demonstrate that multiple tissues in preclinical models are viable for gene signature analysis including WB and lung tissue. Furthermore, they provide robust proof-of-concept that preclinical modeling can provide resolution to assess COR scores as an endpoint for novel TB therapies.

## MATERIALS AND METHODS

### Study design

Using the Gene Expression Omnibus (GEO) repository the following search terms were used: “mus tuberculosis AND Mus musculus” (69 total hits), “macaca tuberculosis” (5 total hits), “oryctolagus tuberculosis” (4 total hits) and “cavia tuberculosis” (1 hit). Within Pubmed, combinations of the following terms were used to search for data sets: tuberculosis, M.tb, biomarkers, RNA, risk scores, preclinical models, mouse, NHP, guinea pig, rabbit, challenge, microarray, sequencing, host gene expression, vaccine, or drug treatment. Data from cell lines, primary sorted cells or single-cell RNA-seq were excluded. We required that included data be from primary reports and not review publications, the data included at least one signature of interest, and groups contained at least two animals per group.

### Microarray dataset pre-processing

All microarray datasets were processed in the R environment (v3.6.1). Normalised microarray datasets were downloaded using the getGEO function of the GEOquery package (*75*) and log_2_-transformed where required. Where datasets were incomplete or uploaded to GEO in an indiscernible format, raw files were downloaded and processed. For Agilent arrays (GSE21149), .txt files were downloaded, and background corrected (Normexp), quantile normalised and filtered for expressed probes using limma (*76*). For Affymatrix arrays (GSE97835), .CEL files were downloaded, and background correction and normalisation was performed using the rma function of the oligo package (*77*).

### RNA-seq dataset pre-processing

Gene counts tables were downloaded using the getGEO function of the GEOquery package. Where required, a normalized counts matrix was generated using DESeq2 (v1.28.1) (*78*) in the R environment (v3.6.1). GSE102440 was processed from raw fastq files since the original assembly used to map the reads was missing a gene required for signature generation. Sample processing from download of .sra files to the generation of a summarised read count matrix was performed on a central shared Linex-based high performance computing service at LSHTM. Sra.toolkit (SRA Toolkit Development Team. SRA Toolkit. https://trace.ncbi.nlm.nih.gov/Traces/sra/sra.cgi?view=software.) was used to download fastq files for BioProject PRJNA397748. STAR (v2.7.3a) (*79*) was used to generate a genome index and subsequently map the reads to the *Macaca mulatta* Mmul_10 assembly. Read counting was performing using the default settings of featureCounts (*80*) specifying a reversely stranded dataset. Quality control steps were performed using FastQC (v0.11.9) (Andrews, S. FastQC: a quality control tool for high throughput sequence data. http://www.bioinformatics.babraham.ac.uk/projects/fastqc/ (2010)) and CollectRnaSeqMetrics from the Picard Tools suite (Broad Institute. Picard Tools. http://broadinstitute.github.io/picard/.). A normalized counts matrix was generated as described above.

### Generation of signature scores

RISK6 (*27*) and Sweeney3 (*34*) signature scores, as well as BAFT2 (*35*) transcript levels, were generated from each dataset as described previously using normalized log2-transformed mean fluorescence intensity values (microarray) or normalized read count values (RNA-seq).

### Statistics

COR signature scores were calculated for each animal and cohorts were compared across conditions using one-way ANOVA with Tukey’s or Dunnetts multiple comparison test correction where there were > 3 groups or evaluations were over time, for comparing two groups at a single timepoint students t-test was used. Significant comparisons are denoted by asterisks in the figure where * = p ≤ 0.05, ** = p ≤ 0.01, *** = p ≤ 0.001, **** = p ≤ 0.0001. Data sets which included animal-matched transcriptional results with time, bacterial burden or pathology endpoint results were evaluated for correlations with Spearman rank-based correlation test and r and p values are reported. Where too few data points were available to meet the requirements of a Spearman correlation, a Pearson correlation was instead used and R^2^ and p values are reported where appropriate.

## Supplementary materials

Fig. S1. Risk signatures correlate strongly with CFU and pulmonary inflammation in an NHP experimental infection model.

Fig. S2. Risk signatures correlate with CFU in a D.O. mouse model.

Table S1. Data captured in searches but excluded based on *a priori* criteria.

## Acknowledgments

The authors would like to thank all authors of the publications used for this reanalysis work. We would also like to thank the VALIDATE network staff for their tireless contributions to supporting TB science and fostering cross continent collaborations.

## Author contributions

Conceptualization: HP, SEL, RNC Methodology: HP, SEL, HFMA, AFG

Investigation: HP, SEL, BDW, HFMA, AFG

Visualization: HP, SEL

Funding acquisition: HP, SEL, HAF, RNC

Project administration: HP, SEL Supervision: SEL, RNC

Writing – original draft: HP, SEL

Writing – review & editing: HP, SEL, BDW, HFMA, SLB, HAF, AFG, RNC

## Data and materials availability

All data are available in the main text or the supplementary materials. This includes a complete list of accession numbers, first author, and preclinical models for all data relating to reanalysis performed in this paper are listed in Tables 3, 4 and 5.

